# Musculoskeletal Model Personalization Affects Metabolic Cost Estimates for Walking

**DOI:** 10.1101/2020.08.05.238857

**Authors:** Marleny Arones, Mohammad Shourijeh, Carolynn Patten, Benjamin J. Fregly

**Affiliations:** Rice University, Department of Mechanical Engineering, Houston, TX, USA; University of California, Department of Physical Medicine and Rehabilitation, Davis, CA, USA

**Keywords:** Metabolic Cost, Model Personalization, Stroke, Gait, EMG-Driven

## Abstract

Assessment of metabolic energy cost as a metric for human performance has expanded across various fields within the scientific, clinical, and engineering communities. As an alternative to measuring metabolic cost experimentally, musculoskeletal models incorporating metabolic cost models have been developed. However, to utilize these models for practical applications, the accuracy of their metabolic cost predictions requires improvement. Previous studies have reported the benefits of using personalized musculoskeletal models for various applications, yet no study has evaluated how model personalization affects metabolic cost estimation. This study investigated the effect of musculoskeletal model personalization on estimates of metabolic cost of transport (CoT) during post-stroke walking using three commonly used metabolic cost models. We analyzed data previously collected from two male stroke survivors with right-sided hemiparesis. The three metabolic cost models were implemented within three musculoskeletal modeling approaches involving different levels of personalization. The first approach used a scaled generic OpenSim model and found muscle activations via static optimization (SOGen). The second approach used a personalized EMG-driven musculoskeletal model with personalized functional axes but found muscle activations via static optimization (SOCal). The third approach used the same personalized EMG-driven model but calculated muscle activations directly from EMG data (EMGCal). For each approach, the muscle activation estimates were used to calculate each subject’s cost of transport (CoT) at different gait speeds using three metabolic cost models (Umberger 2003, Umberger 2010, and Bhargava 2004). The calculated CoT values were compared with published CoT trends as a function of stance time, double support time, step positions, walking speed, and severity of motor impairment (i.e., Fugl-Meyer score). Overall, U10-SOCal, U10-EMGCal, U03-SOCal, and U03-EMGCal were able to produce slopes between CoT and the different measures of walking asymmetry that were statistically similar to those found in the literature. Although model personalization seemed to improve CoT estimates, further tuning of parameters associated with the different metabolic cost models in future studies may allow for realistic CoT predictions. An improvement in CoT predictions may allow researchers to predict human performance, surgical, and rehabilitation outcomes reliably using computational simulations.

## 1 Introduction

Metabolic energy expenditure has been used to evaluate human performance during daily activities such as walking (Waters and Mulroy, 1999; Donelan et al., 2002, 2008; Mian et al., 2006; Long III and Srinivasan, 2013) and athletic activities such as running (Roberts et al., 1998; Chang and Kram, 1999; Teunissen et al., 2007; Franz et al., 2012; Long III and Srinivasan, 2013) and cycling (Davies, 1980; Gnehm et al., 1997; Neptune and Van Den Bogert, 1997; McDaniel et al., 2002; van der Woude et al., 2008). This quantity has also been adopted as a metric to evaluate the design or operational settings of assistive devices (Malcolm et al., 2013; Collins et al., 2015; Galle et al., 2017). Additional applications for which knowledge of metabolic cost is useful include prescription of training intensities (American College of Sports Medicine, 2000), advancement of geriatric medicine (Mian et al., 2006; Canavan et al., 2009; Corbett et al., 2017), treatment of clinical gait disorders (Waters and Mulroy, 1999), and to monitor energy intake and expenditure in obese patients (Brychta et al., 2010). Though various methods exist to measure metabolic cost (Lam and Ravussin, 2016), technical challenges resulting from cost, feasibility, or medical restrictions of the subject population limit the availability of metabolic cost measurements.

With advances in computational biomechanics, musculoskeletal and metabolic cost models have emerged as tools to estimate metabolic energy. These tools have been used in various studies to predict human movement and response to mechanical interventions. Uchida et al. (Uchida et al., 2016b) used the OpenSim simulation framework (Seth et al., 2018) to optimize the design of an assistive device intended to reduce the metabolic cost of running. Dembia et al. (Dembia et al., 2017) simulated an ideal assistive device to minimize the metabolic cost of several individuals walking with heavy loads. Fey et al. (Fey et al., 2012) developed an optimization to identify an optimal prosthetic foot stiffness to minimize metabolic cost for amputee walking. Miller et al. (Miller et al., 2013) used direct collocation optimal control to predict a gait pattern that reduced metabolic cost while minimizing peak axial knee joint contact forces. By predicting the factors, designs, or movements that minimize metabolic energy expenditure, these studies highlight the potential benefits of combining musculoskeletal and metabolic cost models. However, for these predictions to become realistic for practical applications, the accuracy of these combined models needs to improve (Miller, 2014), (Koelewijn et al., 2019).

While published studies that estimated metabolic cost during walking have focused on using scaled generic musculoskeletal models, several studies have reported that personalization of anatomical and physiological characteristics of a musculoskeletal model can influence prediction of muscle forces, joint moments, and novel movements, and these factors also play a role in metabolic cost calculations. Reinbolt et al. (Reinbolt et al., 2007) found that personalization of the functional joint axes is crucial to obtain reliable inverse dynamic joint moments. Studies like (Lloyd and Besier, 2003; Buchanan et al., 2005; Shao et al., 2009; Sartori et al., 2012; Meyer et al., 2017) reported large improvements in joint moment matching when muscle force-generating properties were personalized using an EMG-driven model. Gerus et al. (Gerus et al., 2013) demonstrated that personalizing knee geometry *and* corresponding muscle-tendon parameter values resulted in improved predictions of knee contact force compared to predictions generated by a generic model. Since muscle force estimates depend on joint moments, and the production of muscle forces and activations depends on muscle parameters, the confounding effect of model personalization may also affect metabolic cost estimates. However, to the best of the authors’ knowledge, no study to date has explored this possibility.

This study evaluated the influence of musculoskeletal model personalization on metabolic cost estimates of walking post-stroke. To evaluate the physical realism of different metabolic cost modeling methods, we compared metabolic cost estimates to trends reported in the literature. For individuals post-stroke, Finley and Bastian (Finley and Bastian, 2017) reported a decrease in metabolic cost as walking speed increased and severity of motor impairment decreased. In contrast, for the same subject population, Finley and colleagues (Finley et al., 2015; Finley and Bastian, 2017) also reported that metabolic cost increased as differences in step position, stance time, and double support time increased. The present study evaluated a previously acquired gait data set from which metabolic cost measurements were not available. Our goal was to determine whether any combination of musculoskeletal modeling method and metabolic cost modeling method could reproduce all five physically realistic metabolic cost trends observed in the literature. Since metabolic energy has been adopted as a tool in various scientific, clinical, and engineering fields to evaluate human performance, surgical, and rehabilitation outcomes, the results of this study may be useful for identifying which modeling methods are most likely to predict physically realistic metabolic cost estimates.

## 2 Methods

### 2.1 Experimental Data and Data Processing

Experimental walking data collected from two male stroke survivors – one high functioning and one low functioning - were used as inputs to the metabolic cost analyses (see Table 1 for subject details). Data collected from both subjects included marker-based motion capture, ground reaction, and surface and fine-wire electromyographic (EMG) data (Table 2, 16 channels per leg). The data were collected at different walking speeds using a split-belt instrumented treadmill with belts tied. Walking speeds ranged from slower than self-selected to a maximum comfortable speed. For the high functioning subject, the speed range was from 0.4 to 0.8 m/s in increments of 0.1 m/s, while for the low functioning subject, the speed range was from 0.35 to 0.65 m/s also in increments of 0.1 m/s. Additional experimental data were also collected for static and isolated joint movement trials. A static standing trial was collected for model scaling purposes. Isolated joint motion trials were collected for each hip, knee, and ankle for purposes of personalizing the model’s lower body joint axes. All functional axes for the joint of interest were exercised during each isolated joint motion trial (Reinbolt et al., 2005, 2008).

**Table 1.** Clinical characteristics of study participants.

| Gender | Age | Paretic Limb | Lower Limb Fugl-Meyer | Self Selected Speed (m/s) |
| --- | --- | --- | --- | --- |
| M | 79 | R | 32 | 0.5 |
| M | 62 | R | 25 | 0.35 |

**Table 2.**
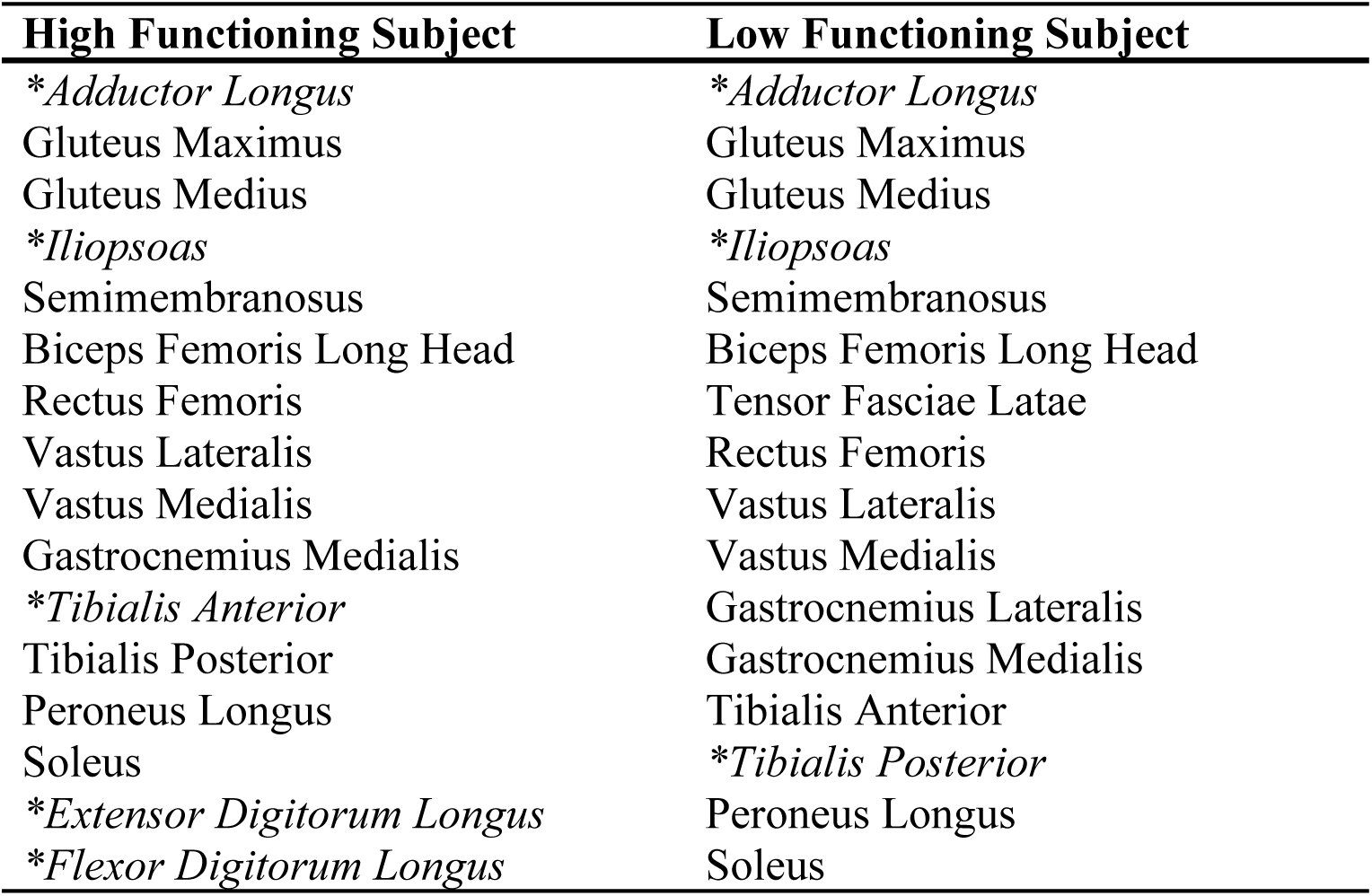
A list of muscle groups from which surface or fine-wire (*) EMG signal was recorded.

| High Functioning Subject | Low Functioning Subject |
| --- | --- |
| *\*Adductor Longus* | *\*Adductor Longus* |
| Gluteus Maximus | Gluteus Maximus |
| Gluteus Medius | Gluteus Medius |
| *\*Iliopsoas* | *\*Iliopsoas* |
| Semimembranosus | Semimembranosus |
| Biceps Femoris Long Head | Biceps Femoris Long Head |
| Rectus Femoris | Tensor Fasciae Latae |
| Vastus Lateralis | Rectus Femoris |
| Vastus Medialis | Vastus Lateralis |
| Gastrocnemius Medialis | Vastus Medialis |
| *\*Tibialis Anterior* | Gastrocnemius Lateralis |
| Tibialis Posterior | Gastrocnemius Medialis |
| Peroneus Longus | Tibialis Anterior |
| Soleus | *\*Tibialis Posterior* |
| *\*Extensor Digitorum Longus* | Peroneus Longus |
| *\*Flexor Digitorum Longus* | Soleus |

The ground reaction and marker motion data were low-pass filtered using a fourth-order zero-phase lag Butterworth filter with a cut-off frequency of 7/*tf* Hz (Hug, 2011), where *tf* is the period of the gait cycle being processed (Meyer et al., 2017). On average, this variable cut-off frequency caused data collected at a normal walking speed to be filtered at approximately 6 Hz.

The EMG data were high-pass filtered (40 Hz), demeaned, rectified, and low-pass filtered (3.5/*tf* Hz) using a fourth-order zero-phase lag Butterworth filter. EMG amplitudes for each muscle were normalized to the maximum value over all trials and resampled to 101 time points per gait cycle, as described in (Meyer et al., 2017).

### 2.2 Musculoskeletal Model

A generic full-body OpenSim musculoskeletal model (Rajagopal et al., 2016; Seth et al., 2018) served as the starting point for all three metabolic cost analyses. The generic model used for both subjects started with 40 Hill-type muscle-tendon actuators per leg and 37 degrees of freedom (DOFs), including 3 DOF hip joints, 1 DOF knee joints, 2 DOF ankle joints. For the high functioning subject, six muscles without related EMG data were eliminated (extensor hallucis longus, flexor hallucis longus, gracilis, piriformis, sartorius, tensor fascia latae). For the low functioning subject, seven muscles without related EMG data were eliminated (extensor digitorum longus, flexor digitorum longus, extensor hallucis longus, flexor hallucis longus, gracilis, piriformis, sartorius). The remaining muscles actuated hip flexion-extension, hip adduction-abduction, hip internal-external rotation, knee flexion-extension, ankle flexion-extension, and ankle inversion–eversion on each leg.

### 2.3 Joint Model Personalization

Personalization of the joint functional axes for the hip, knee, and ankle on each leg was performed by following a two-step process. First, the geometry of the generic OpenSim model was scaled to match the dimensions of each subject using OpenSim’s Scale tool and the static standing trial. Second, marker positions and functional axes of the model’s lower body’s joints were personalized as described in (Reinbolt et al., 2005, 2008; Meyer et al., 2016; Sauder et al., 2019). The personalization process involved using nonlinear optimization to adjust the positions and orientations of the model’s lower body joints and marker triads placed on the body segments. The cost function minimized the sum of squares of errors between the experimental and model marker positions from all isolated joint motion trials and one walking trial analyzed together. The optimization process was performed using Matlab’s *lsqnonlin* algorithm, which iteratively ran OpenSim’s inverse kinematics analyses to calculate marker location errors.

### 2.4 Muscle-tendon Model and Geometry Personalization

Experimental data from ten gait trials collected at each available speed were used to calibrate an EMG-driven model for each subject. Before performing EMG-driven model calibration, we analyzed marker data from each gait trial using OpenSim’s Inverse Kinematics tool to generate joint angle trajectories. OpenSim’s Inverse Dynamics tool was then used to calculate the joint moments to be produced by muscle forces. Next, a surrogate model of each subject’s musculoskeletal geometry was generated to allow the EMG-driven model to modify muscle geometry (Meyer et al., 2016, 2017). The surrogate model was developed by sampling a wide range of joint angle combinations for the lower limbs using a Latin hypercube design. The muscle-tendon lengths and moment arms for each muscle were then calculated using OpenSim’s Muscle Analysis tool. Linear regression using least squares was used to fit muscle-tendon lengths as a polynomial function of the corresponding joint angles actuated by each muscle. Muscle-tendon velocity was then defined as the first derivative of the muscle-tendon length polynomial with respect to time while muscle moment arms were defined as the negative of the first derivative of the muscle-tendon length polynomial with respect to corresponding joint angles (An et al., 1984).

To personalize the model’s muscle-tendon force-generating properties, we allowed our EMG-driven model calibration process to modify three types of parameters: EMG-to-activation parameters (electromechanical delays, activation dynamics time constants, activation non-linearization shape factors, and EMG scale factors), Hill-type muscle-tendon model parameters (optimal muscle fiber lengths, and tendon slack lengths), and surrogate musculoskeletal geometry parameters. With the use of Matlab’s *fmincon* algorithm using sequential quadratic programming, the model parameter values were adjusted to best match calculated experimental inverse dynamic moments and published passive joint moments (Silder et al., 2007). Additionally, penalty terms were added to the cost function to discourage substantial divergence of model parameters from their original values. A detailed description of our EMG-driven modeling approach can be found in Meyer et al. (Meyer et al., 2017).

### 2.5 Analysis

We developed three musculoskeletal modeling approaches, each derived from the same generic OpenSim model and implemented in Matlab, to evaluate the extent to which model personalization affects estimated metabolic cost. The first approach used a scaled generic OpenSim model where muscle activations were calculated via static optimization using quadratic programming (SO-Gen). The second approach used the subject’s personalized EMG-driven model but with muscle activations found via static optimization (SO-Cal). The third approach used the same personalized EMG-driven model but found muscle activations using the EMG-driven model (EMG-Cal) (Figure 1). The muscle activations found by each approach were used to calculate each subject’s cost of transport (CoT in J/m/kg) for different gait speeds using metabolic cost models published by Umberger (Umberger et al., 2003) (U03), Umberger et al. (Umberger, 2010) (U10), and Bhargava et al. (Bhargava et al., 2004) (B04).

**Figure 1.**
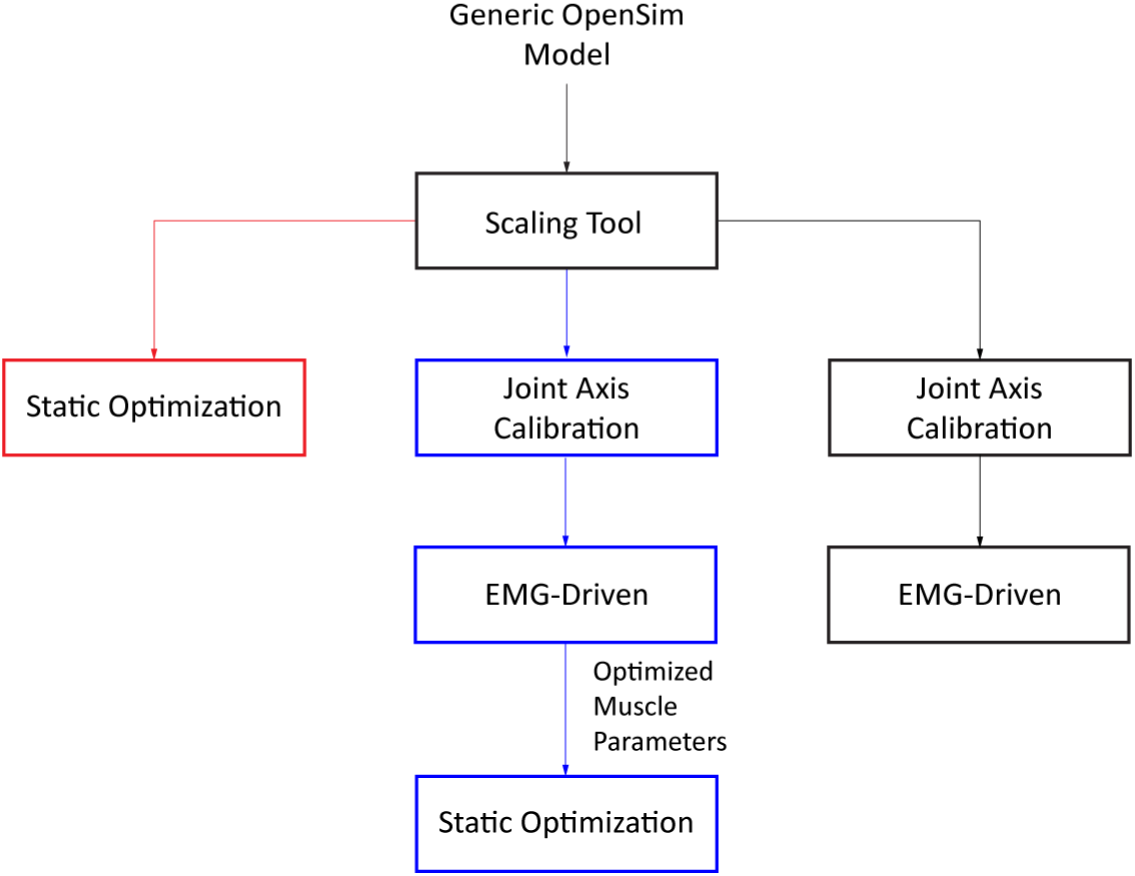
Flowchart of the three different approaches used to obtain estimates of muscle activations (SOGen: red, SOCal: blue, EMGCal: black).

To evaluate the physical realism of each musculoskeletal modeling approach/metabolic cost model combination, we identified five experimental trends in the literature for how CoT varies with other quantities for individuals post-stroke. The first three quantities were step position asymmetry, stance time asymmetry, and double-support time asymmetry that were reported to increase as the cost of transport increased (Finley and Bastian, 2017). Inversely, the two other quantities were speed and Fugl-Meyer score that were observed to decrease as the cost of transport increased. Differences or asymmetries in step positions, stance time, and double-support time were calculated as specified in (Finley et al., 2015; Finley and Bastian, 2017). The relationship between estimated CoT and the variables of interest was compared against the linear correlations reported in (Finley and Bastian, 2017). Thus, a t-test analysis, comparing the slopes produced between CoT estimates and variables of interest against those found in the literature, was performed for each modeling approach and metabolic cost model. The t-test analysis was performed for all correlations except CoT and Fugl-Meyer score due to the limitation of only two subjects.

To investigate the effect of either increasing or decreasing muscle specific tension away from its nominal value of 61 N/cm (Meyer et al., 2017), we iteratively ran static optimization with varying muscle specific tension values using the personalized model for the high functioning patient. All iterations of this static optimization were performed at a fixed gait speed of 0.5 m/s (self-selected speed).

## 3 Results

When the nine modeling combinations were used to predict the positive or negative correlations of CoT with other quantities as reported in (Finley and Bastian, 2017), U10-SOCal, U10-EMGCal, U03SO-Cal, and U03-EMGCal predicted statistically similar slopes for all four variables (excluding Fugl-Meyer score) (*p* ≥ 0.05) (Table 3) (Figures 2-5). In addition, all modeling combinations showed a decrease in CoT as Fugl-Meyer score increased (Table 4-5).

**Table 3.** Resulting p-values obtained from a t-test analysis between the calculated and experimental CoT estimates. Statistically similar slopes are in bold.

|  | Umberger 2003 |  |  | Umberger 2010 |  |  | Bhargava 2004 |  |  |
| --- | --- | --- | --- | --- | --- | --- | --- | --- | --- |
|  | SOGen | SOCal | EMGCal | SOGen | SOCal | EMGCal | SOGen | SOCal | EMGCal |
| Speed | 0.033 | **0.12** | **0.23** | 0.033 | **0.13** | **0.20** | 0.0064 | 0.0089 | 0.025 |
| Step Position Contribution | **0.056** | **0.22** | **0.52** | **0.30** | **0.21** | **0.40** | 0.0062 | 0.020 | **0.20** |
| Stance Time Differences | **0.26** | **0.30** | **0.39** | **0.24** | **0.29** | **0.34** | **0.22** | **0.25** | **0.34** |
| Double Support Time Differences | **0.088** | **0.20** | **0.45** | **0.064** | **0.19** | **0.37** | 0.014 | 0.023 | **0.22** |

**Table 4.** Cost of transport estimates for the three metabolic cost models and all three personalization methods across Fugl-Meyer scores for the high functioning subject.

|  | Umberger 2003 |  |  | Umberger 2010 |  |  | Bhargava 2004 |  |  |
| --- | --- | --- | --- | --- | --- | --- | --- | --- | --- |
|  | SOGen | SOCal | EMGCal | SOGen | SOCal | EMGCal | SOGen | SOCal | EMGCal |
| Max CoT | 4.66 | 6.24 | 6.58 | 6.17 | 7.10 | 8.24 | 2.72 | 2.45 | 2.35 |
| Min Cot | 7.05 | 10.98 | 11.35 | 8.69 | 11.96 | 13.14 | 3.29 | 3.67 | 3.50 |
| Mean CoT | 5.56 | 8.13 | 8.73 | 7.10 | 9.04 | 10.46 | 2.89 | 2.89 | 2.92 |

**Table 5.** Cost of transport estimates for the three metabolic cost models and all three personalization methods across Fugl-Meyer scores for the low functioning subject.

|  | Umberger 2003 |  |  | Umberger 2010 |  |  | Bhargava 2004 |  |  |
| --- | --- | --- | --- | --- | --- | --- | --- | --- | --- |
|  | SOGen | SOCal | EMGCal | SOGen | SOCal | EMGCal | SOGen | SOCal | EMGCal |
| Max CoT | 4.68 | 6.74 | 7.74 | 5.91 | 7.55 | 9.23 | 2.37 | 2.62 | 3.25 |
| Min Cot | 7.65 | 11.63 | 12.79 | 9.21 | 12.57 | 14.33 | 3.76 | 4.11 | 4.51 |
| Mean CoT | 5.98 | 8.74 | 9.82 | 7.38 | 9.60 | 11.30 | 3.03 | 3.19 | 3.77 |

**Figure 2.**
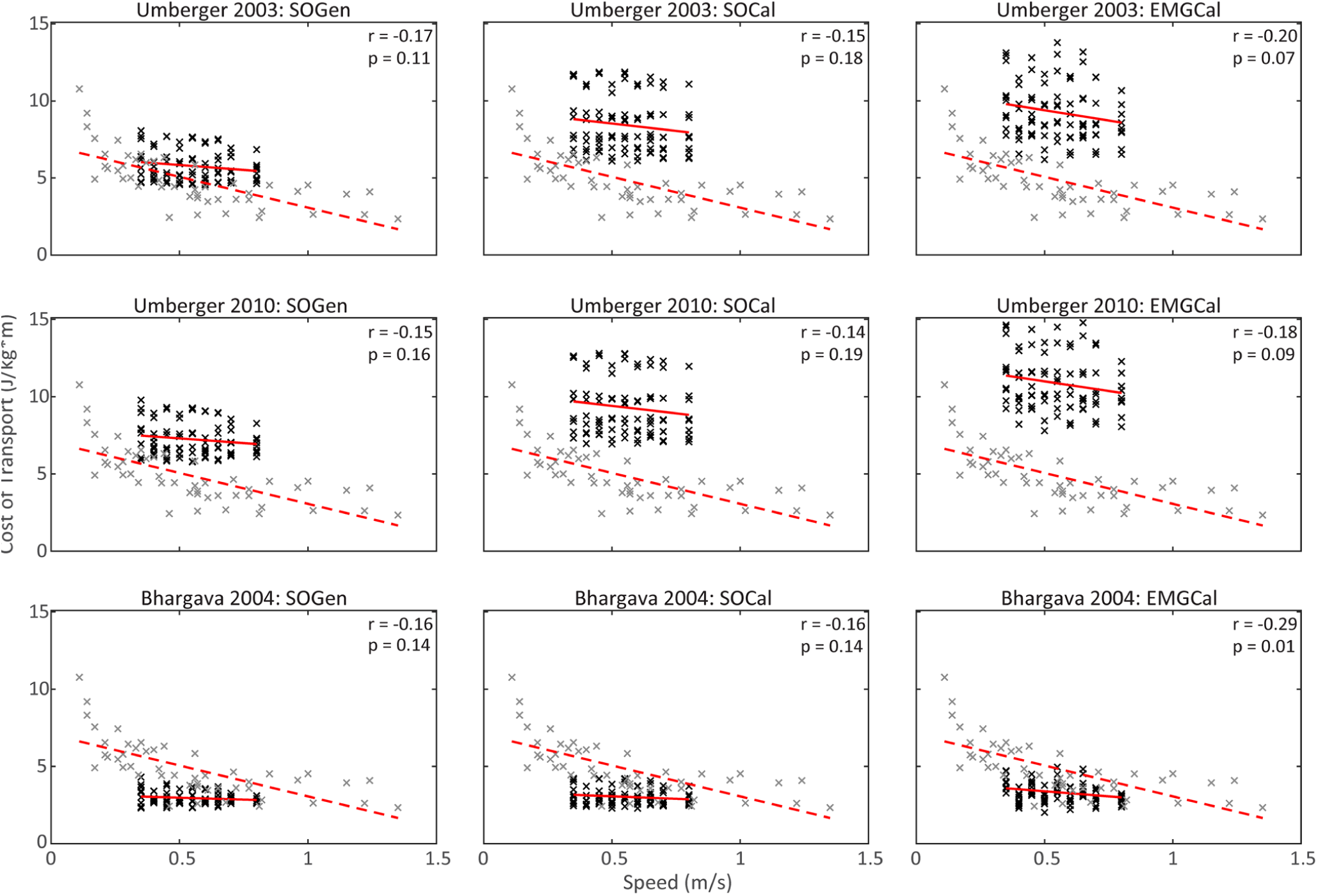
Cost of transport estimates for the three metabolic cost models and for all three personalization methods (black) across speeds. Experimental data reported in Finley and Bastian (Finley and Bastian, 2017) are shown in grey.

**Figure 3.**
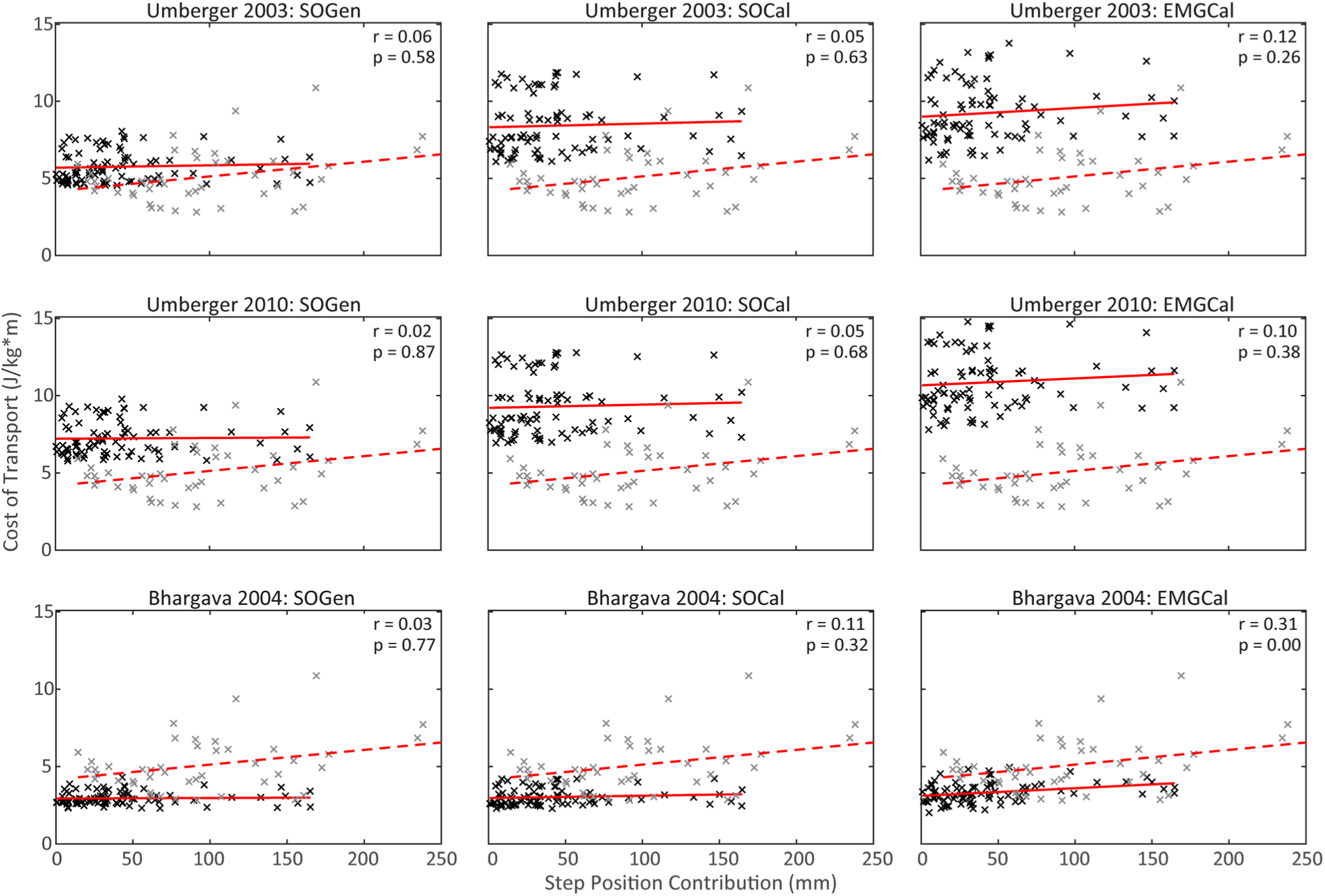
Cost of transport estimates for the three metabolic cost models and for all three personalization methods (black) across step position contribution to step length differences. Experimental data reported in Finley and Bastian 2017 are shown in grey.

**Figure 4.**
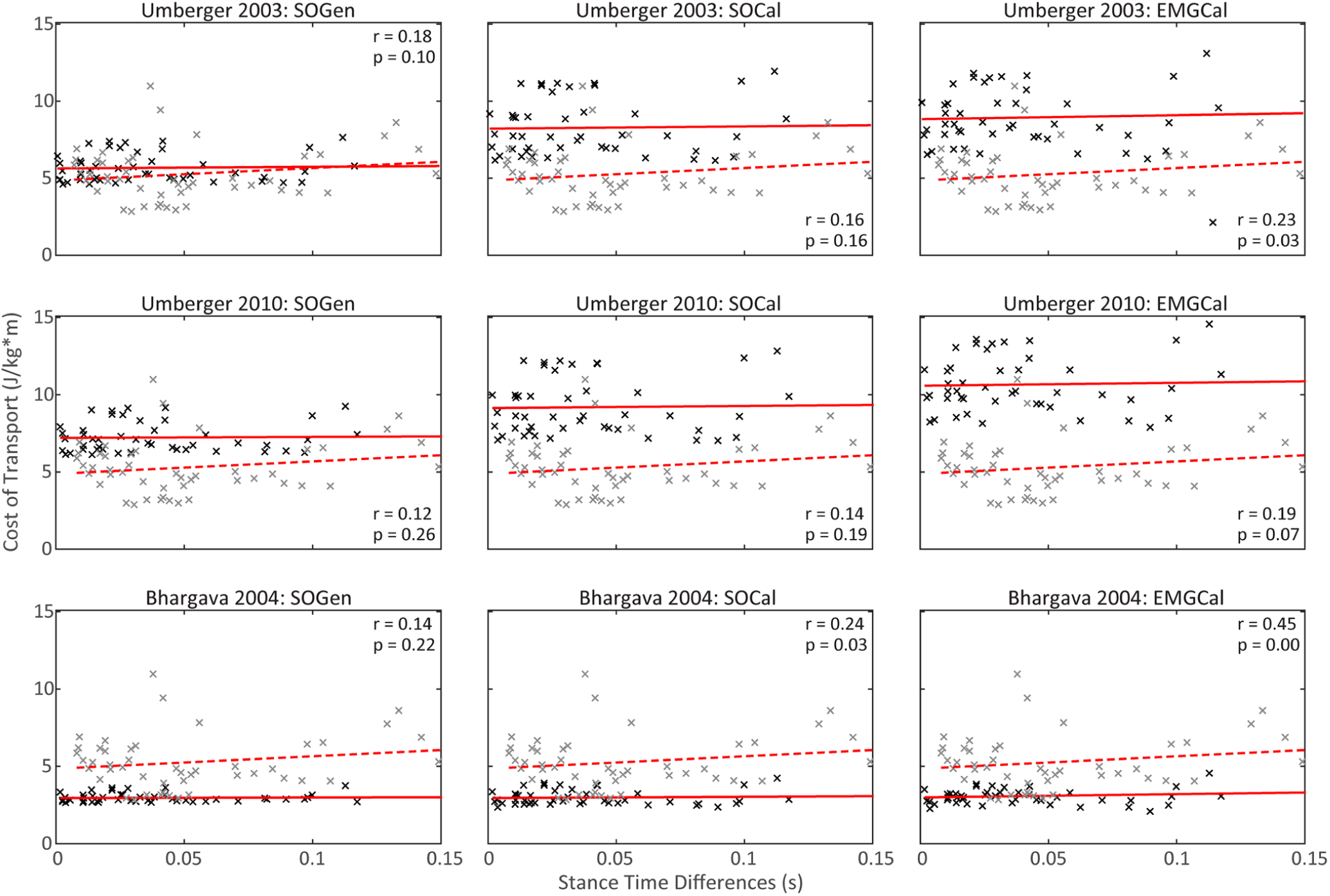
Cost of transport estimates for the three metabolic cost models and for all three personalization methods across stance time differences. Experimental data reported in Finley and Bastian 2017 are shown in grey.

**Figure 5.**
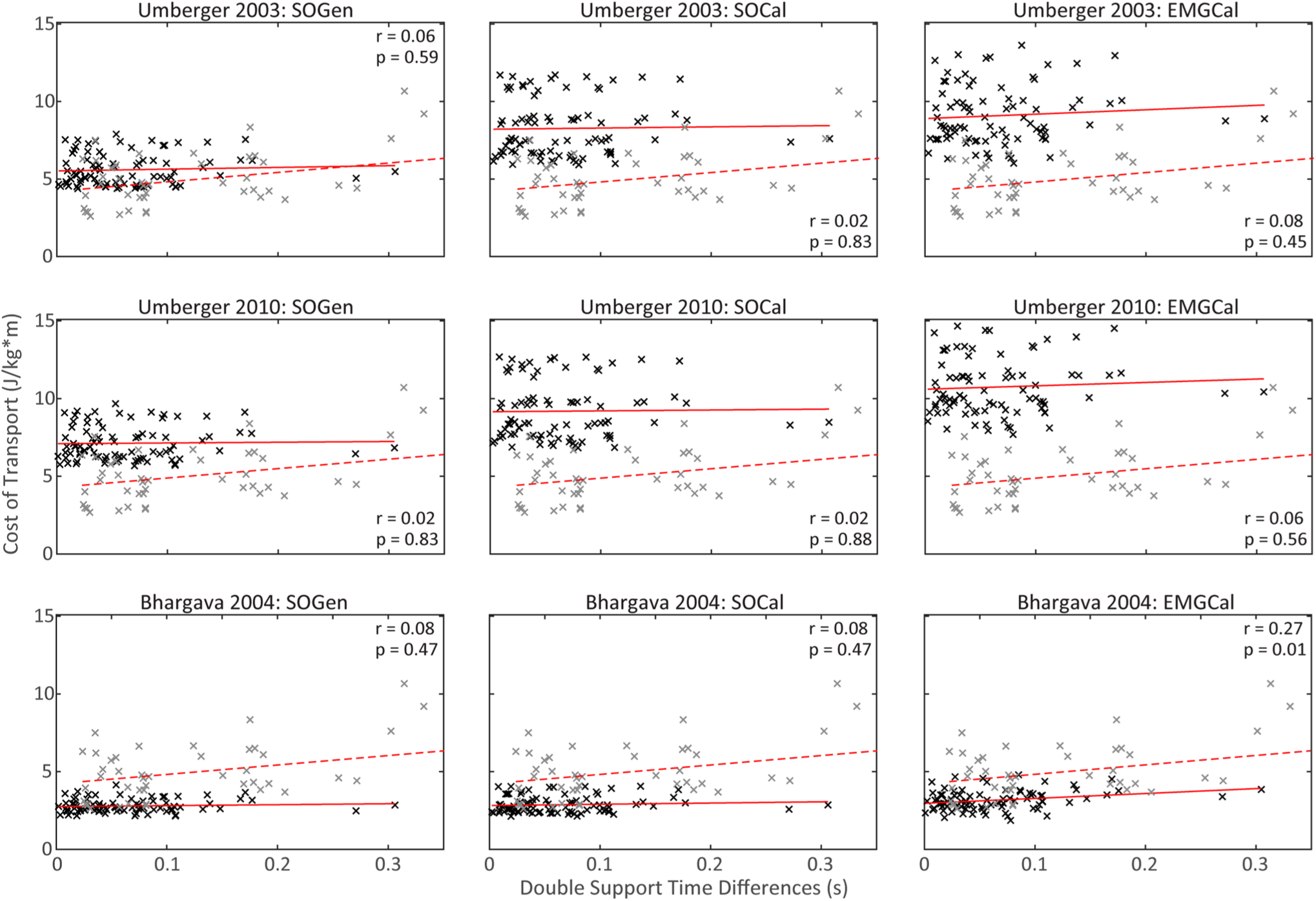
Cost of transport estimates for the three metabolic cost models and for all three personalization methods across double support time differences. Experimental data reported in Finley and Bastian 2017 are shown in grey.

An inverse relationship was observed between changes in CoT and changes in muscle specific tension (Figure 6). To compensate for CoT overestimation by both Umberger models (U03 and U10), muscle specific tension had to be increased to 75-135 N/cm^2^ for U03 and 90-135 N/cm^2^ for U10. With this change, the resulting CoT estimates fit within one standard deviation of the average reported by Finley and Bastian, 2017. Inversely, to compensate for CoT underestimation by the Bhargava model, muscle specific tension had to be decreased to 37 N/cm^2^ for the resulting CoT estimates to lie within the range of CoT values published by (Miller, 2014; Koelewijn et al., 2019).

**Figure 6.**
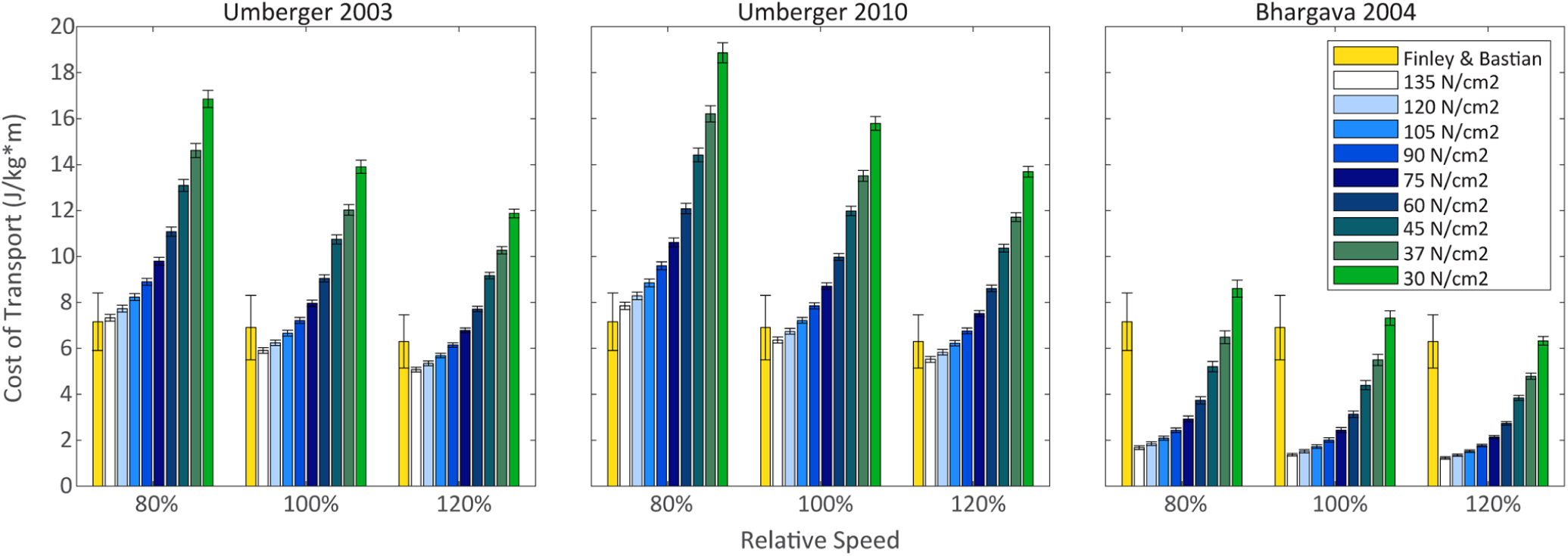
Increase in cost of transport as the specific muscle strength decreases using the SO-Cal modeling combination for the high functioning patient across three speeds (relative to self-selected speed – 100%).

## 4 Discussion

This study evaluated the effect of musculoskeletal model personalization on metabolic cost estimates for walking obtained using three common metabolic cost models: Umberger 2003 (U03), Umberger *et al.* 2010 (U10), and Bhargava et al. 2004 (B04). These three models were implemented within three musculoskeletal modeling approaches: scaled generic musculoskeletal models with muscle activations found by static optimization (SOGen) (Shourijeh and Fregly, 2020), calibrated EMG-driven musculoskeletal models with muscle activations found by static optimization (SOCal), and calibrated EMG-driven musculoskeletal models with muscle activations found from experimental EMG data (EMGCal). These nine combinations were investigated using previously published walking data collected from two individuals post-stroke, and correlations between estimated CoT and walking speed, step position asymmetry, step length asymmetry, stance time asymmetry, double support time asymmetry, and Fugl-Meyer score were compared with published data (Finley and Bastian, 2017). All nine modeling combinations predicted the correct positive and negative correlations between CoT and the five selected quantities as observed in a post-stroke population. However, only the personalized models (SOCal andEMGCal) paired with Umberger 2003 and Umberger 2010 produced statistically similar slopes for all four quantities in comparison to the linear models found in the literature. These findings suggest that a calibrated EMG-driven musculoskeletal model combined with either one of Umberger’s models improved CoT estimates.

In general, only SOGen predicted CoT estimates that were within one standard deviation of the published experimental average for all three metabolic cost models. EMGCal predicted CoT estimates that were larger than those obtained from both SOGen and SOCal for all three metabolic cost models. Additionally, the level of model personalization did not affect the tendency of Bhargava’s model to underestimate CoT nor the tendency of both Umberger’s models to overestimate CoT (Figure 2). A similar pattern was reported in Miller (Miller, 2014), which analyzed the performance of B04 and U10, among other models. However, in our study, the combination of U03 or U10 used with EMGCal produced CoT values that were almost twice as large as published CoT averages. Studies such as Ong (Ong et al., 2019), which used a model based on U03, and Miller (Miller, 2014) which used U10, reported CoT values within the range of 2 to 8 J/Kg*m, whereas U10-EMG resulted in CoT values ranging from 7 to 15 J/Kg*m. This magnitude difference was not expected but may be due to minimization of a metabolic cost term within the optimization cost function used for estimating muscle activations in both studies. The observation of minimization of metabolic cost may explain why CoT magnitudes produced by SOGen were closest to those reported in (Finley and Bastian, 2017) for all three metabolic cost models. Additionally, in contrast to the studies mentioned above, our study also used calibrated EMG-driven models to calculate CoT. The absence of muscle activation minimization may help explain why our EMGCal paired with any metabolic model produced CoT estimates that were larger than those produced by SOGen and SOCal.

Although all modeling combinations produced the positive or negative trends relative to published data, not all of them resulted in slopes that were statistically similar to those found in the literature. This finding is similar to a recent study by Koelwijn et al. (Koelewijn et al., 2019), where several metabolic cost models (including U03 and B04) were shown to predict the correct trends for CoT as a function of walking slope or gait speed in accordance to trends found in the literature (Margaria, 1968). Additionally, Koelwijn et al. (Koelewijn et al., 2019) found that B04 tended to underestimate CoT values in comparison to experimental values, which is in agreement with our findings.

Although future work can be directed toward optimizing parameters within the metabolic cost models, we tested how changing muscle specific tension affected the magnitude of the predicted CoT values. As reported in (Russell, Esposito and Miller, 2018), decreasing the muscle specific tension increased metabolic cost and vice versa. Bhargava’s model needed muscle specific tension to be around 37 N/cm^2^ rather than 61 N/cm^2^ to match experimental averages the best. This new value is closer to values used within OpenSim (Delp et al., 2007) and studies such as (Miller, 2014; Koelewijn et al., 2019). Future work to optimize muscle specific tension could improve the magnitude of CoT predictions (Hamner and Delp, 2013; Uchida et al., 2016a; Falisse et al., 2019).

One of the major limitations of this study was the absence of experimental metabolic cost data for evaluating our model-based estimates. This limitation makes it difficult to determine the accuracy of our modeling methods. Furthermore, none of the musculoskeletal models used in our analysis included muscles to actuate the upper body; therefore, the metabolic cost for all nine cases may be an underestimation. However, since Finley and Bastian’s dataset were collected from subjects who were allowed to use handrails, their measurements may be underestimates as well. Additionally, we were limited to two subjects with the extensive EMG datasets required to perform the study, which could have impacted our statistical results.

In conclusion, this study investigated the effect of musculoskeletal model personalization on estimated metabolic cost for post-stroke walking as obtained from three commonly used metabolic cost models and three musculoskeletal model personalization methods. Previously collected walking data from two stroke survivors were used to analyze correlations between estimated CoT and various variables commonly associated with gait asymmetry. Although all modeling combinations exhibited the correct positive and negative correlations observed in (Finley and Bastian, 2017), U10-SOCal, U10-EMGCal, U03-SOCal, and U03-EMGCal produced statistically similar slopes to those found in the literature for all gait asymmetry variables (excluding Fugl-Meyer score). While the estimated CoT values were not close to reference CoT averages (Finley and Bastian, 2017), we observed that additional tuning of muscle specific tension easily changed the CoT estimates. Therefore, future tuning of muscle specific tension along with other parameters associated with the metabolic cost models may further improve CoT estimates. Our results suggest the use of a personalized EMG-driven model to estimate muscle forces paired with either one of Umberger’s energy expenditure models to help the prediction of CoT estimates during walking for individuals post-stroke. Since metabolic energy has been adopted as a tool in various scientific, clinical, and engineering fields to evaluate performance, surgical, and rehabilitation outcomes, the results of this study may help improve the reliability of metabolic cost predictions generated using musculoskeletal models.

## 6 Author Contributions

CP and BJF performed experiments; MA performed all model personalization tasks, prepared figures, and drafted manuscript; MA and MS analyzed data; MA, MS, and BF interpreted results of experiments; MA, MS, CP and BF revised manuscript.

## 9 Supplementary Material

Additional figures can be found in Supplementary Material

