## Supplementary Material for "Musculoskeletal Model Personalization Affects Metabolic Cost Estimates for Walking" (from another version of this preprint: version 1, 2020-08-06)

### 1 Supplementary Tables

**Table 4.** Average RMS errors between muscle produced joint moments from SO-Gen and EMG-Driven in comparison to SO-Cal across all trials

| Joint Moments | High Functioning Subject | | | | | | Low Functioning Subject | | | | | |
|---|---|---|---|---|---|---|---|---|---|---|---|---|
| | Paretic Leg | | | Non-paretic Leg | | | Paretic Leg | | | Non-paretic Leg | | |
| | SOGen | SOCal | EMGCal | SOGen | SOCal | EMGCal | SOGen | SOCal | EMGCal | SOGen | SOCal | EMGCal |
| Hip flexion | 2.29 | 0 | 5.12 | 3.60 | 0 | 6.65 | 1.00 | 0 | 7.03 | 0.71 | 0 | 5.46 |
| Hip adduction | 2.85 | 0 | 5.53 | 2.04 | 0 | 7.31 | 0.88 | 0 | 4.83 | 1.03 | 0 | 5.97 |
| Hip rotation | 0.42 | 0 | 2.00 | 0.49 | 0 | 2.60 | 0.78 | 0 | 1.74 | 0.43 | 0 | 2.17 |
| Knee flexion | 1.57 | 0 | 5.15 | 2.20 | 0 | 5.97 | 9.49 | 0 | 7.68 | 1.75 | 0 | 4.75 |
| Ankle rotation | 0.36 | 0 | 6.47 | 0.70 | 0 | 5.73 | 2.60 | 0 | 4.33 | 0.84 | 0 | 6.18 |
| Ankle eversion | 1.18 | 0 | 1.86 | 0.49 | 0 | 2.91 | 2.17 | 0 | 1.15 | 1.02 | 0 | 3.31 |

**Table 5.** Average RMS errors between muscle activations produced from static optimization to those obtained from the EMG-Driven model.

| Muscle | High Functioning Subject | | | | | | Low Functioning Subject | | | | | |
|---|---|---|---|---|---|---|---|---|---|---|---|---|
| | Paretic Leg | | | Non-paretic Leg | | | Paretic Leg | | | Non-paretic Leg | | |
| | SOGen | SOCal | EMGCal | SOGen | SOCal | EMGCal | SOGen | SOCal | EMGCal | SOGen | SOCal | EMGCal |
| Addbrev | 0.02 | 0.02 | 0 | 0.02 | 0.02 | 0 | 0.03 | 0.01 | 0 | 0.02 | 0.02 | 0 |
| Addlong | 0.03 | 0.02 | 0 | 0.06 | 0.02 | 0 | 0.05 | 0.02 | 0 | 0.05 | 0.02 | 0 |
| AddmagDist | 0.02 | 0.02 | 0 | 0.02 | 0.01 | 0 | 0.02 | 0.01 | 0 | 0.02 | 0.02 | 0 |
| AddmadIsch | 0.02 | 0.02 | 0 | 0.01 | 0.04 | 0 | 0.02 | 0.01 | 0 | 0.02 | 0.02 | 0 |
| AddmagMid | 0.02 | 0.02 | 0 | 0.02 | 0.02 | 0 | 0.02 | 0.01 | 0 | 0.02 | 0.02 | 0 |
| AddmagProx | 0.16 | 0.16 | 0 | 0.02 | 0.02 | 0 | 0.14 | 0.15 | 0 | 0.06 | 0.04 | 0 |
| Bflh | 0.23 | 0.21 | 0 | 0.18 | 0.11 | 0 | 0.02 | 0.02 | 0 | 0.25 | 0.23 | 0 |
| Bsfh | 0.33 | 0.24 | 0 | 0.26 | 0.07 | 0 | 0.29 | 0.07 | 0 | 0.41 | 0.30 | 0 |
| Tfl | - | - | - | - | - | - | 0.45 | 0.38 | 0 | 0.35 | 0.46 | 0 |
| Edl | 0.24 | 0.03 | 0 | 0.19 | 0.07 | 0 | - | - | - | - | - | - |

| | | | | | | | | | | | | |
|---|---|---|---|---|---|---|---|---|---|---|---|---|
| Fdl | 0.01 | 0.02 | 0 | 0.42 | 0.39 | 0 | - | - | - | - | - | - |
| Gaslat | 0.09 | 0.03 | 0 | 0.43 | 0.31 | 0 | 0.04 | 0.03 | 0 | 0.09 | 0.08 | 0 |
| Gasmed | 0.17 | 0.09 | 0 | 0.11 | 0.04 | 0 | 0.21 | 0.08 | 0 | 0.20 | 0.13 | 0 |
| Glmax1 | 0.04 | 0.14 | 0 | 0.05 | 0.08 | 0 | 0.04 | 0.05 | 0 | 0.07 | 0.12 | 0 |
| Glmax2 | 0.08 | 0.13 | 0 | 0.09 | 0.15 | 0 | 0.06 | 0.05 | 0 | 0.07 | 0.15 | 0 |
| Glmax3 | 0.02 | 0.04 | 0 | 0.02 | 0.02 | 0 | 0.02 | 0.02 | 0 | 0.03 | 0.04 | 0 |
| Glmed1 | 0.23 | 0.28 | 0 | 0.32 | 0.11 | 0 | 0.13 | 0.12 | 0 | 0.14 | 0.09 | 0 |
| Glmed2 | 0.14 | 0.21 | 0 | 0.07 | 0.06 | 0 | 0.13 | 0.11 | 0 | 0.13 | 0.24 | 0 |
| Glmed3 | 0.24 | 0.27 | 0 | 0.15 | 0.14 | 0 | 0.25 | 0.11 | 0 | 0.24 | 0.21 | 0 |
| Glmin1 | 0.10 | 0.04 | 0 | 0.17 | 0.09 | 0 | 0.02 | 0.03 | 0 | 0.08 | 0.14 | 0 |
| Glmin2 | 0.11 | 0.04 | 0 | 0.09 | 0.12 | 0 | 0.02 | 0.06 | 0 | 0.08 | 0.12 | 0 |
| Glmin3 | 0.08 | 0.06 | 0 | 0.07 | 0.04 | 0 | 0.03 | 0.11 | 0 | 0.15 | 0.16 | 0 |
| Iliacus | 0.18 | 0.05 | 0 | 0.28 | 0.07 | 0 | 0.16 | 0.04 | 0 | 0.22 | 0.19 | 0 |
| Perbrev | 0.08 | 0.02 | 0 | 0.08 | 0.04 | 0 | 0.07 | 0.04 | 0 | 0.14 | 0.11 | 0 |
| Perlong | 0.13 | 0.04 | 0 | 0.15 | 0.08 | 0 | 0.13 | 0.02 | 0 | 0.17 | 0.09 | 0 |
| Psoas | 0.17 | 0.03 | 0 | 0.20 | 0.03 | 0 | 0.39 | 0.10 | 0 | 0.40 | 0.42 | 0 |
| Recfem | 0.16 | 0.12 | 0 | 0.26 | 0.20 | 0 | 0.17 | 0.09 | 0 | 0.20 | 0.25 | 0 |
| Semimem | 0.22 | 0.22 | 0 | 0.21 | 0.14 | 0 | 0.10 | 0.09 | 0 | 0.05 | 0.06 | 0 |
| Semiten | 0.22 | 0.15 | 0 | 0.18 | 0.09 | 0 | 0.10 | 0.09 | 0 | 0.23 | 0.19 | 0 |
| Soleus | 0.38 | 0.25 | 0 | 0.08 | 0.07 | 0 | 0.02 | 0.06 | 0 | 0.49 | 0.30 | 0 |
| Tibant | 0.23 | 0.32 | 0 | 0.26 | 0.12 | 0 | 0.45 | 0.06 | 0 | 0.32 | 0.20 | 0 |
| Tibpost | 0.03 | 0.04 | 0 | 0.09 | 0.16 | 0 | 0.32 | 0.34 | 0 | 0.26 | 0.16 | 0 |
| Vasint | 0.13 | 0.11 | 0 | 0.07 | 0.05 | 0 | 0.02 | 0.05 | 0 | 0.12 | 0.06 | 0 |
| Vaslat | 0.12 | 0.05 | 0 | 0.14 | 0.08 | 0 | 0.13 | 0.11 | 0 | 0.11 | 0.05 | 0 |
| Vasmed | 0.15 | 0.09 | 0 | 0.04 | 0.03 | 0 | 0.02 | 0.02 | 0 | 0.10 | 0.04 | 0 |

### 2 Muscle Models

**Legend:**
$\tilde{l}_M$– Normalized Muscle Length
$\tilde{F}_P$ – Passive Force
$\tilde{v}_M$– Normalized Muscle Velocity
$\tilde{F}_A$– Active Force

**Force Length Model:**
$b11 = 0.814483478343008;$
$b21 = 1.055033428970575;$
$b31 = 0.162384573599574;$
$b41 = 0.063303448465465;$
$b12 = 0.433004984392647;$
$b22 = 0.716775413397760;$
$b32 = -0.029947116970696;$
$b42 = 0.200356847296188;$
$b13 = 0.1;$
$b23 = 1;$
$b33 = 0.5 * \sqrt{0.5};$
$b43 = 0;$

$$\tilde{F}_A = b11 e^{\frac{-0.5*(\tilde{l}_M - \mathrm{b21})^2}{(b31 + b41\tilde{l}_M)^2}} + b12 e^{\frac{-0.5*(\tilde{l}_M - \mathrm{b22})^2}{(b32 + b42\tilde{l}_M)^2}} + b13 e^{\frac{-0.5*(\tilde{l}_M - \mathrm{b23})^2}{(b33 + b43\tilde{l}_M)^2}}$$

**Force Velocity Model:**
$d1 = -8.665833008200368;$
$d2 = 7.623504943764548;$
$d3 = 3.565407127548163;$
$d4 = 0.968466940650055;$
$d5 = -0.394453556647392;$
$d6 = 8.169548497993912;$

$$\tilde{F}_A = d1 + d2 * \mathrm{atan}(\mathrm{d3} + \mathrm{d4} * \mathrm{atan}(\mathrm{d5} + \mathrm{d6} * \tilde{v}_M));$$

**Passive Force Model:**
$c1 = 3.384513491001194;$
$c2 = -74.090638944399615;$
$c3 = -2.443789456896774;$

$$\tilde{F}_P = c_1 e^{c_2\left(e^{\tilde{l}_M}\right)^{c_3}}$$
